# Accounting for Uncertainty in Gene Tree Estimation: Summary-Coalescent Species Tree Inference in a Challenging Radiation of Australian Lizards

**DOI:** 10.1101/056085

**Authors:** Mozes P.K. Blom, Jason G. Bragg, Sally Potter, Craig Moritz

## Abstract

Accurate gene tree inference is an important aspect of species tree estimation in a summary-coalescent framework. Yet, in empirical studies, inferred gene trees differ in accuracy due to stochastic variation in phylogenetic signal between targeted loci. Empiricists should therefore examine the consistency of species tree inference, while accounting for the observed heterogeneity in gene tree resolution of phylogenomic datasets. Here, we assess the impact of gene tree estimation error on summary-coalescent species tree inference by screening ~2000 exonic loci based on gene tree resolution prior to phylogenetic inference. We focus on a phylogenetically challenging radiation of Australian lizards (genus *Cryptoblepharus*, Scincidae) and explore effects on topology and support. We identify a well-supported topology based on all loci and find that a relatively small number of high-resolution gene trees can be sufficient to converge on the same topology. Adding gene trees with decreasing resolution produced a generally consistent topology, and increased confidence for specific bipartitions that were poorly supported when using a small number of informative loci. This corroborates coalescent-based simulation studies that have highlighted the need for a large number of loci to confidently resolve challenging relationships and refutes the notion that low-resolution gene trees introduce phylogenetic noise. Further, our study also highlights the value of quantifying changes in nodal support across locus subsets of increasing size (but decreasing gene tree resolution). Such detailed analyses can reveal anomalous fluctuations in support at some nodes, suggesting the possibility of model violation. By characterizing the heterogeneity in phylogenetic signal among loci, we can account for uncertainty in gene tree estimation and assess its effect on the consistency of the species tree estimate. We suggest that the evaluation of gene tree resolution should be incorporated in the analysis of empirical phylogenomic datasets. This will ultimately increase our confidence in species tree estimation using summary-coalescent methods and enable us to exploit genomic data for phylogenetic inference.

---

With the development of novel sequencing methods, phylogenomic datasets are generated that can contain hundreds to thousands of orthologous loci (Blair and Murphy 2011; Yang and Rannala 2012; McCormack et al. 2013). This steep change in the availability of genetic markers has led to a focus on phylogenetic methods that accommodate the frequently observed incongruence in evolutionary histories among loci (Jennings and Edwards 2005; Edwards 2009; Lemmon and Lemmon 2013; Nater et al. 2015). Topological discordance due to biological sources such as incomplete lineage sorting, can be expected under a wide variety of realistic evolutionary scenarios and therefore needs to be evaluated when inferring species trees (Pamilo and Nei 1988; Maddison 1997; Degnan and Rosenberg 2009). Most species tree methods aim to account for this heterogeneity in coalescent histories by incorporating the multi-species coalescent (MSC) model (Knowles 2009; Liu et al. 2009; Degnan and Rosenberg 2009; Nakhleh 2013; Liu et al. 2015a). Using the MSC and independent genetic markers, these methods weigh distinct species tree hypotheses by comparing the observed distribution in gene trees with an expected distribution given a species tree hypothesis. Full-coalescent sequence based methods that jointly infer gene trees and species tree, such as *BEAST (Heled and Drummond 2010) or BEST (Liu 2008), are preferable but also remain computationally intractable with a large number of loci (Leaché and Rannala 2011). Thus, the use of such methods remains unfeasible for most phylogenomic studies and alternative approaches have been developed that alleviate computational burden (Kubatko et al. 2009; Liu et al. 2010; Mirarab et al. 2014b).

Coalescent methods based on summary statistics (‘summary-coalescent’), use a twoAstep approach where individual gene trees are initially inferred and species tree inference is conducted by summarizing across the resulting collection of gene trees. summary-coalescent methods are computationally cheap and have become increasingly popular as the species tree method of choice for empirical phylogenomic studies using full-sequence data (Lemmon et al. 2012; Ilves and López-Fernández 2014; Bond et al. 2014; Leaché et al. 2014; Giarla and Esselstyn 2015). However, summary-coalescent methods have been criticized for disregarding uncertainty in gene tree estimation and their unrestricted adoption has been questioned (Gatesy and Springer 2013; 2014; Springer and Gatesy 2016). Although statistically inconsistent (Roch and Steel 2015), concatenating loci can under some circumstances result in a better estimate of the underlying species tree than a summary-coalescent tree that is based on gene trees with poor phylogenetic signal (Mirarab et al. 2014a). The potential problems associated with gene tree accuracy has motivated some phylogeneticists to favor concatenation over summary-coalescent approaches (Gatesy and Springer 2013; 2014; Springer and Gatesy 2016; but see Edwards et al. 2016). However, the relative importance of gene tree inaccuracy remains undetermined (Huang et al. 2010), since other studies have suggested that accurate species trees can still be inferred even in the presence of gene tree estimation error (Roch and Warnow 2015). Thus, empirical studies incorporating summary-coalescent methods should validate the consistency of species tree inference, while accounting for the potential uncertainty in gene tree estimation.

There are two main sources underlying gene tree estimation error. First, model misspecification can result in systematic error, where alternative gene trees are consistently recovered due to the fit of erroneous models of sequence evolution (Jeffroy et al. 2006; Kumar et al. 2012; Doyle et al. 2015). Second, if the phylogenetic information content (PIC) of an individual locus is relatively low, the inferred gene tree will be inaccurate and conflicting topologies are equally likely (‘low resolution’). The distribution of PIC in current phylogenomic studies is often uneven, since genetic markers frequently vary in length and/or mutation rate (Faircloth et al. 2012; Lemmon et al. 2012; Lanier and Knowles 2012). The inferred gene trees in empirical studies will therefore differ in precision and gene tree estimation error can be regarded as a stochastic artifact inherent to the method used for generating phylogenomic sequence data, the loci targeted and the success of reassembling contigs.

Whereas the performance of species tree methods has been extensively tested (McCormack et al. 2009; Leaché and Rannala 2011; Mirarab et al. 2014a), these simulations often do not address the observed heterogeneity in PIC among loci and the corresponding variation in resolution of inferred gene trees. Simulation studies that have explicitly accounted for variation in PIC show that the use of genes with higher mutation rates (Huang et al. 2010; Lanier et al. 2014; Giarla and Esselstyn 2015) or longer length (Liu et al. 2015b) can significantly increase the accuracy of species tree estimation. Yet, the effect of adding low-resolution loci to an informative dataset remains unclear; some simulations suggest that such gene trees do not affect the species tree estimate (Lanier et al. 2014), whereas others report a decrease in performance rather than an improvement in accuracy (Liu et al. 2015b).

The concerns regarding gene tree estimation error and the observed variation of gene tree resolution in empirical datasets have motivated us to explore the performance of summary-coalescent species tree inference while explicitly considering the resolution of the included gene trees. Recently developed target-capture methods provide opportunity to generate largeAscale DNA sequence datasets (Faircloth et al. 2012; Lemmon et al. 2012; Bragg et al. 2015; Jones and Good 2016) and to use empirical data for characterizing the effect of gene tree resolution on species tree inference. Here we thoroughly explore an exon-capture dataset (~2000 loci) to address this issue and simultaneously aim to resolve a challenging radiation of Australian lizards.

Lizards of the genus *Cryptoblepharus* Wiegmann (Reptilia: Squamata: Scincidae) are small scansorial skinks that range through three broad, geographic regions: The Ethiopian-Malagasy (southwest Indian Ocean), Indo-Pacific and Australian (Ineich and Blanc 1988; Rocha et al. 2006; Hayashi et al. 2009; Blom 2015a). A thorough revision of 396 Australian *Cryptoblepharus* individuals (Horner and Adams 2007), using 45 genetic (allozyme) and 33 morphological markers, increased the number of described Australian species from seven to 25. Based on genetic differences between taxa, the diversification of Australian lineages has likely occurred in two discrete radiations (clades A and B). The crown age for each clade is around 5 Ma., but clade B contains more lineages (11 and 17 taxa respectively) and so has diversified more rapidly (Blom et al. 2016). Phylogenetic relationships in recent, rapid radiations can be notoriously difficult to resolve due to the widespread presence of incomplete lineage sorting and the potential reliance on genetic markers that evolve slowly relative to the rate of speciation (Giarla and Esselstyn 2015). Hence, this Australian group of skinks provides an excellent opportunity to study potential sensitivities in the inference process, while resolving radiations with distinct rates of diversification (i.e. clades A and B).

Here we quantify the impact of stochastic gene tree estimation error on summary-coalescent species tree inference by screening loci based on gene tree resolution prior to species tree estimation. We use a recently developed measure (Salichos and Rokas 2013; Salichos et al. 2014) to quantify the degree of conflict among all bipartitions present in a set of bootstrapped trees as a proxy for gene tree resolution. It is not the aim of this study to explicitly compare the performance between species tree methods (Leache and Rannala 2011), investigate the effects of missing data (Streicher et al. 2016) or explore the effect of systematic gene tree estimation error (Doyle et al. 2015). Rather we aim to characterize the heterogeneity in gene tree resolution that is frequently observed in empirical datasets and to quantify its effect on topology and support of species trees. Applying such thorough evaluations will enable empiricists to benefit from the additional value that novel molecular approaches offer to the field of phylogenetics.

## Materials and Methods

### Taxon Sampling

Based on results from Horner and Adams (2007), we selected a single representative for each of the 28 identified allozyme lineages of Australian *Cryptoblepharus* (Supplementary Table 1). Three of these lineages have not been diagnosed as species, due to morphological and geographic overlap with currently described species (Horner 2007). Here they are treated as separate taxa because the available samples exhibit a considerable degree of genetic differentiation from other species (Horner and Adams 2007).

Most specimens included in this study are held in the collections of the Museum and Art Gallery of the Northern Territory (MAGNT), Western Australian Museum (WAM) or Queensland Museum (QM) and were used in the initial taxonomic revision. We used tissues available from the Australian Biological Tissue Collection at the South Australian Museum, unless tissues previously analyzed were depleted. For these species, we used recently collected field samples. Museum specimens from Horner and Adams (2007) have associated allozyme profiles, but we screened recently collected samples for morphological characteristics and sequenced a mitochondrial marker (*ND2*) to verify correct species assignment. We did not verify lineage assignment for one species, *C. gurrmul*, since the amount of available tissue was very limited. *C. gurrmul* is endemic to a small number of islands and is the only *Cryptoblepharus* species present in the sampled location (Horner 2007). We chose another taxon within the Eugongylus group (*Bassiana duppereyi*) as an outgroup, based on existing phylogenetic hypotheses (Brandley et al. 2015).

### Exon Capture Design, Library Preparation and Sequencing

The design of the exon-capture kit used in this study is outlined in detail in Bragg et al. (2015). Briefly, we identified a set of potential exon targets with a balanced base composition, in the *Anolis* genome and identified their orthologs in transcriptomes of three species from genera related to *Cryptoblepharus* (*Carlia rubrigularis, Lampropholis coggeri* and *Saproscincus basiliscus;* Singhal 2013). A total of 3320 loci were targeted (>200 base pairs), with a total target length of 4.31 Mb (including a representative of each exon from each of the three species). Based on these exon targets, Roche NimbleGen designed and synthesized a SeqCap EZ Developer Library as our probe set. These probes capture homologous targets with high efficiency across the entire Eugongylus group, to which *Cryptoblepharus* belongs (Bragg et al. 2015).

We extracted genomic DNA from liver tissue stored either frozen or in RNALater, following the slating-out method of Sunnucks and Hales (1996). We prepared genomic libraries with ~1400 ng. input DNA per sample and according to the protocol of Meyer and Kircher (2010), using modifications of Bi et al. (2012). In brief, library preparation consisted of blunt-end repair, adapter ligation, adapter fill-in and was followed by two independent index-PCRs to reduce PCR bias. Each sample had a unique barcode for pooling DNA for the hybridization. We assessed DNA concentrations using a Nanodrop (Thermo scientific) and the distribution of fragment lengths on 1.5% agarose gels. Barcoded libraries were pooled in equimolar ratios prior to hybridization. The exon-capture hybridization was performed following SeqCap EZ Developer Library user’s guide (Roche NimbleGen). We assessed the quality of the hybridizations using qPCR following methods of Bi et al. (2012). The qPCR assays used specific primers to assess enrichment of targeted regions, and de-enrichment of non-targeted regions, of the genome. In addition, the quantity and quality of the hybridizations were measured using a Bioanalyzer (Agilent Technologies), to quantify the concentration of the pre- and post-capture libraries. Once the libraries passed all aforementioned quality checks (i.e. successful enrichment), they were submitted for sequencing (Functional Genomics Lab QB3 core facility, UC Berkeley). We sequenced the enriched libraries (100bp pairedAend) on a single Illumina HiSeq 2000 lane.

### Read Processing and Assembly

Illumina sequencing reads were processed using a workflow that was described previously by Singhal (2013), and is available at https://github.com/MVZSEQ. The workflow removes duplicate, low complexity and contaminant reads, and trims adaptors and low quality bases (TRIMMOMATIC v0.22, Bolger et al. 2014). Overlapping reads were merged using FLASH (v1.2.2, Magoc and Salzberg 2011).

Cleaned sequencing reads were assembled using an approach described by Bragg et al. (2015). Briefly, for each sample, each locus was assembled separately, after identifying reads with homology to the encoded protein (using blastx, v2.2.25, expectation value = 1EA9, Altschul et al. 1990). These reads were then assembled with VELVET (K values 31, 41, 51, 61, 71 and 81; v1.2.08, Zerbino and Birney 2008). Contigs for a locus were combined using CAP3 (parameter values -o 20 -p 99, version date 08/06/13, Huang and Madan 1999), and trimmed to the exon boundaries using EXONERATE (v2.2.0, Slater and Birney 2005). Where multiple contigs were assembled, we used a reciprocal best blastx hit criterion to select a contig orthologous to the targeted *Anolis* exon (see Bragg et al. 2015).

### Alignment and Data Pre-processing for Species Tree Inference

High-quality sequence alignments are essential for correct phylogenetic inference (Zwickl et al. 2014) but visual inspection of alignment quality, as traditionally conducted, is challenging with thousands of loci (Lemmon and Lemmon 2013; Blom 2015a). We developed a flexible bioinformatic workflow for alignment and alignment filtering of exonic sequences, EAPhy (v0.9, Blom 2015b), and generated high-quality alignments for each specific analysis. In brief, EAPhy aligned sequences using MUSCLE (v3.8.31, Edgar 2004), performed checks to ensure coding of amino acids and removed missing data from the ends of the alignments. We visually inspected alignments after filtering if individual sequences still deviated significantly (more than three aminoAacids in a seven base window) from the alignment consensus or contained more than one stop codon. EAPhy generates the desired input format (i.e. Phylip, Fasta) for most species tree methods and the complete pipeline can be found at (https://github.com/MozesBlom/EAPhy).

### Assessing Stochastic Gene Tree Estimation Error

Stochastic gene tree estimation error is driven by a lack of phylogenetic signal, and the phylogenetic informativeness of loci is often characterized as the total number or relative ratio of parsimony informative sites (e.g. Leache et al. 2014). However, the relationship between gene tree accuracy and the number of informative sites is not necessarily proportional. For example, only a small proportion of sites might prove informative for challenging clades with high rates of diversification because most informative sites will only differentiate between major clades or in- and outgroup taxa (Townsend 2007). Thus, whereas the number of informative sites might be high for a given locus, the gene tree accuracy in some parts of the tree might still be relatively poor. An alternative way to evaluate gene tree estimation is to compare the consistency of bootstrap replicates. This could be calculated as the average bootstrap support (BS) value across all internal nodes. This would provide an indication of the overall consistency, but does not necessarily estimate the accuracy of the inferred gene tree across bootstrap replicates. For example, if a certain bipartition is observed in 51/100 bootstrapped trees, it is unknown whether the second most common, but conflicting, bipartition is supported by the remaining 49 bootstrapped trees or only a few. Here we assume that the estimate of each bipartition is more accurate when the second most common, conflicting, bipartition is observed at low frequency. To estimate the accuracy of each gene tree, we used the ‘Tree Certainty’ (TC) score introduced by Salichos et al. (2013; 2014). The TC score is the sum of ‘Internode Certainty’ estimates, which represent the support for each bipartition of a given topology by considering its frequency in a set of bootstrapped trees, jointly with that of the most prevalent conflicting bipartition in the same set of trees. If gene tree estimation is precise, most bipartitions across the tree are consistently recovered in much higher frequencies than conflicting bipartitions and TC, the sum of internode certainty estimates, is large. Alternatively, TC will be small if many internodes have a low resolution, suggesting that a high frequency of conflicting bipartitions have been inferred across bootstrap replicates. For each locus, we used RAxML (v8.1, Stamatakis 2014) to infer the gene tree with the highest likelihood out of 10 replicates (GTR + Γ) and estimated the TC score based on 100 bootstrap replicates. We calculated the TC score by inferring the majority rule consensus tree across bootstrap replicates and summing the internode certainty scores for each of the inferred bipartitions of the consensus tree. Finally, to assess a potential correlation between locus length and TC score, we used a linear model in R (‘lm’ function, v3.1.2, R Core Team 2014).

### Species Tree Inference

Prior to estimating species trees with coalescent based methods, we inferred a maximum-likelihood (ML) phylogeny based on a concatenated alignment of all loci. When loci are concatenated, variation in genealogical histories is not considered explicitly, and this can result in inflated support metrics (Kubatko and Degnan 2007; Edwards et al. 2007; Knowles 2009; Roch and Steel 2015). However, in addition to providing an initial phylogenetic hypothesis, we generated a concatenated ML phylogeny to test whether the inferred branch lengths were predictive for the degree of discordance among loci. That is, we expected that a shorter interval between species splits would result in a higher degree of incomplete lineage sorting and thus require more loci to confidently infer the underlying species history in a coalescent-based framework (Degnan and Rosenberg 2009). Lastly, we assessed whether species tree methods that incorporate a Full-coalescent model benefit from the identification of loci with a high TC-score. Full-coalescent based methods are computationally intractable with large numbers of loci and the ranking of loci by TC score can prioritize the inclusion of more informative loci over others, potentially resulting in more accurate inference than when using a random subset.

*Concatenated ML species tree*.— We generated a dataset where all *Cryptoblepharus* species were represented and where each alignment had a minimum length of a 150 bp. We then used PartitionFinder (rcluster search, v1.1.1, Lanfear et al. 2012) to identify the optimal partitioning scheme and substitution model for the concatenated alignment by considering both gene and codon position. Using the optimal partitioning strategy and appropriate substitution model (GTR + Γ), we inferred a ML phylogeny from the concatenated dataset using RAxML. The tree with the highest likelihood score was selected out of 100 replicate searches and BS values calculated for each bipartition based on 1000 bootstrap replicates.

*Summary-coalescent species trees*.— To evaluate the impact of adding loci with decreasing phylogenetic resolution, we used the RAxML gene tree and TC score for each locus, and inferred a summary-coalescent species tree for subsets of loci using ASTRAL II v.4.7.6 (Fig. 1; Mirarab and Warnow 2015). We used ASTRAL II since it performs better or equally well in comparison to other summary-coalescent methods (Chou et al. 2015; Ogilvie et al. 2016), is computationally efficient and uses unrooted gene trees.

**Figure 1.**
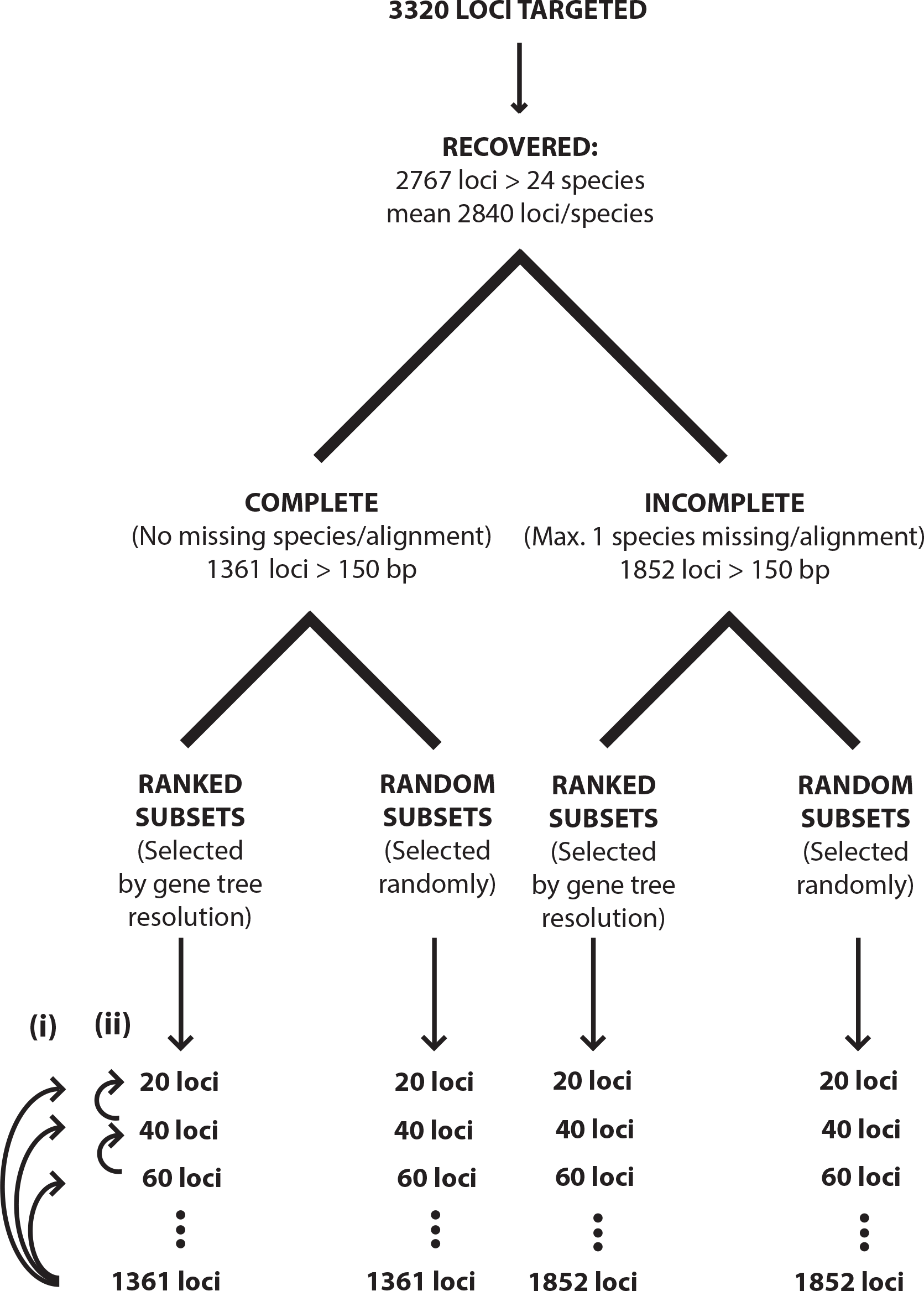
Schematic of the data structure used for analyses. Analyses were conducted on a dataset that contained alignments without missing species and a dataset that also included alignments that missed up to one species. Each dataset was divided in subsets, containing gene trees that were either picked randomly or ranked by Tree Certainty (TC) scores. After inferring an ASTRAL species tree for each subset of loci, Robinson-Foulds distances were calculated by comparing the species tree for each locus subset with the species tree based on all loci (i) or the species tree based on the previous locus subset (ii).

Locus subsets of equal size were either chosen randomly from all available loci (‘random’) or by highest remaining TC value (‘ranked’). For the first 200 loci, we iteratively increased the size of each subset with 20 additional loci. During subsequent iterations, the size of each subset was increased with 100 loci until all available loci were included (Fig. 1). We inferred a summary-coalescent species tree during each iteration for both the random and ranked locus subsets and used multi-locus bootstrapping, as incorporated in ASTRAL II, to calculate BS values for each bipartition (100 replicates).

We inferred species trees for complete gene trees where all species were present (Fig. 1; ‘complete’ dataset) and for incomplete gene trees where up to one of the 28 species could be absent (Fig. 1; ‘incomplete’ dataset). It is possible to include incomplete gene trees since ASTRAL II uses all quartet trees supported in each gene tree for scoring the species tree, whether the taxa are all present or not (S. Mirarab, pers. comm.). We set the criterion of one missing species for two reasons. First, we calculated TC scores for gene trees where data for up to three species were missing (i.e. ~ 10%), and there was not a substantial improvement in the number of alignments with high TC scores when allowing for two or three missing sequences. Second, we have been conservative regarding the degree of missing data permitted since the heuristics of ASTRAL II, which account for the uncertainty that is potentially introduced by tolerating missing taxa, seem to generate consistent results with various degrees of missing data (Xi et al. 2016) but have not been tested exhaustively.

For the bipartitions that were supported in both the concatenated ML phylogeny and the summary-coalescent species trees, we extracted the internode branch length (in mean number of nucleotide substitutions per site) defining each bipartition in the ML phylogeny and calculated the corresponding average BS value across summary-coalescent trees. To calculate the average BS values for each supported bipartition, we took the summary-coalescent trees based on the alignments from the incomplete dataset and used species trees based on ranked subsets of loci. The size of these locus subsets increased iteratively with 100 loci in the range of 100 to 1800 loci. We assessed a potential correlation between branch length and average BS for each bipartition using a linear model in R (‘lm’ function), but excluded the bipartitions that are strongly supported (BS = 100) across all subsets since these are uninformative to the model.

*Full-coalescent species trees*.— We ranked loci by TC score and generated subsets of the 20, 30, 40 and 50 highest ranked loci (complete dataset - i.e. no missing taxa). Furthermore, we generated eight random subsets of 20 loci and five random subsets of 30 loci, from the 200 loci with the highest TC score. We then used *BEAST v2.1.3. (Bouckaert et al. 2014) to infer both gene and species trees. We ran each *BEAST run in duplicate, with separate starting seeds, using a GTR + Γ substitution model with four r rate categories, a strict clock model and applied a birth-death species tree prior. Each analysis was run for 200 million generations and we sampled the MCMC chain every 200,000 generations. We discarded the first 10% as burn-in, used Tracer v1.5 to check for convergence (estimated sample size; ESS > 200) and LogCombiner v2.1.3. to combine the posterior sample of trees across runs. *BEAST analyses that failed to converge were rerun with 400 million generations (required for two random subsets of 30 loci). Since subsets of loci with the 40 and 50 highest ranked TC scores still failed to converge, we only report the inferred species trees based on 20 and 30 loci. The species tree for each subset was individually summarized using TreeAnnotator v2.1.3.

### Evaluating the Effect of Gene Tree Resolution

With ASTRAL II, we inferred a summary-coalescent species tree for each locus subset. This resulted in a sequence of species trees, where each tree is inferred based on a larger (ranked or random) subset of loci (see Fig. 1). To evaluate potential changes in topology across locus subsets of increasing size, we compared species tree topologies using a Robinson-Foulds (RF) tree distance calculation (Robinson and Foulds 1981). The RF distance represents the number of bipartition changes between two trees. By comparing the species tree for each locus subset with the species tree based on all loci for the complete or incomplete dataset, we quantified a) the number of changes in topology when adding gene trees randomly or b) the number of changes in topology when adding ranked gene trees with decreasing resolution. Since the ‘true’ species tree was unknown (in comparison to typical simulation studies), we compared the species tree inferred for each locus subset to the tree based on all loci, assuming that the species tree based on ‘complete evidence’ was the best species tree given the dataset. In addition, we also compared the species tree for each subset of loci to the species tree based on the previous subset of loci (e.g. subset with 300 loci vs. subset with 200 loci), to circumvent the inevitability of increasing topological concordance as the number of loci approaches the full dataset (see Fig. 1). Lastly, as for the ASTRAL comparisons, we evaluated topological changes across *BEAST runs by comparing the species tree for each subset to the ASTRAL species tree based on all loci (incomplete dataset - i.e. 1852 loci, one individual missing).

To calculate RF distances between trees, we rooted each species tree (both ASTRAL and *BEAST) on the *B. duppereyi* outgroup and used the *’symmetric_difference’* function in the Python module Dendropy v.4.0.2 (Sukumaran and Holder 2010). All analyses were conducted for both the complete and incomplete dataset.

## RESULTS AND DISCUSSION

### Characteristics of Loci

We used a custom exon-capture system that was designed for related Eugongylus skinks, and observed considerable capture success across most species of *Cryptoblepharus*. On average, 2840 out of 3320 targeted loci were successfully assembled for each individual, with a mean coverage of 140X (Supplementary Table 1). These results are in line with Bragg et al. (2015), who achieved similar capture success across lizard genera with up to ~40 million years of divergence.

The majority of loci were sequenced across many species (i.e. 2767 loci-24 species or more; Supplementary Fig. S1), but a significant proportion of the loci (~60%) was not recovered across all species in the study. In total, 1484 loci were successfully recovered across all target species and 1361 of these were longer than 150 bp (complete dataset; mean length 384 bp/alignment). If we relaxed the criterion for missing data and allowed up to one species missing per alignment, an additional 491 loci (each > 150 bp) were included, yielding a total dataset of 1852 genes (incomplete dataset; mean length 410 bp/alignment).

We found substantial variation in TC scores for both the complete and incomplete dataset (Fig. 2), illustrating the heterogeneity of resolution among gene trees. This motivated us to further examine how such variation might affect species tree inference. In comparison to the complete dataset, the incomplete dataset yielded more loci and the average gene tree resolution increased for each similar sized locus subset ranked by TC score (Fig. 2). To verify whether allowing for additional missing species (n = 2 and 3 missing) would result in an even further increase in average TC score per locus subset, we also calculated the TC scores for genes that had up to three species missing per alignment. This did not result in a sizeable gain in gene tree resolution across similar sized locus subsets (Fig. 2) and we therefore limited our analyses to the complete and incomplete (up to one species missing) dataset.

**Figure 2.**
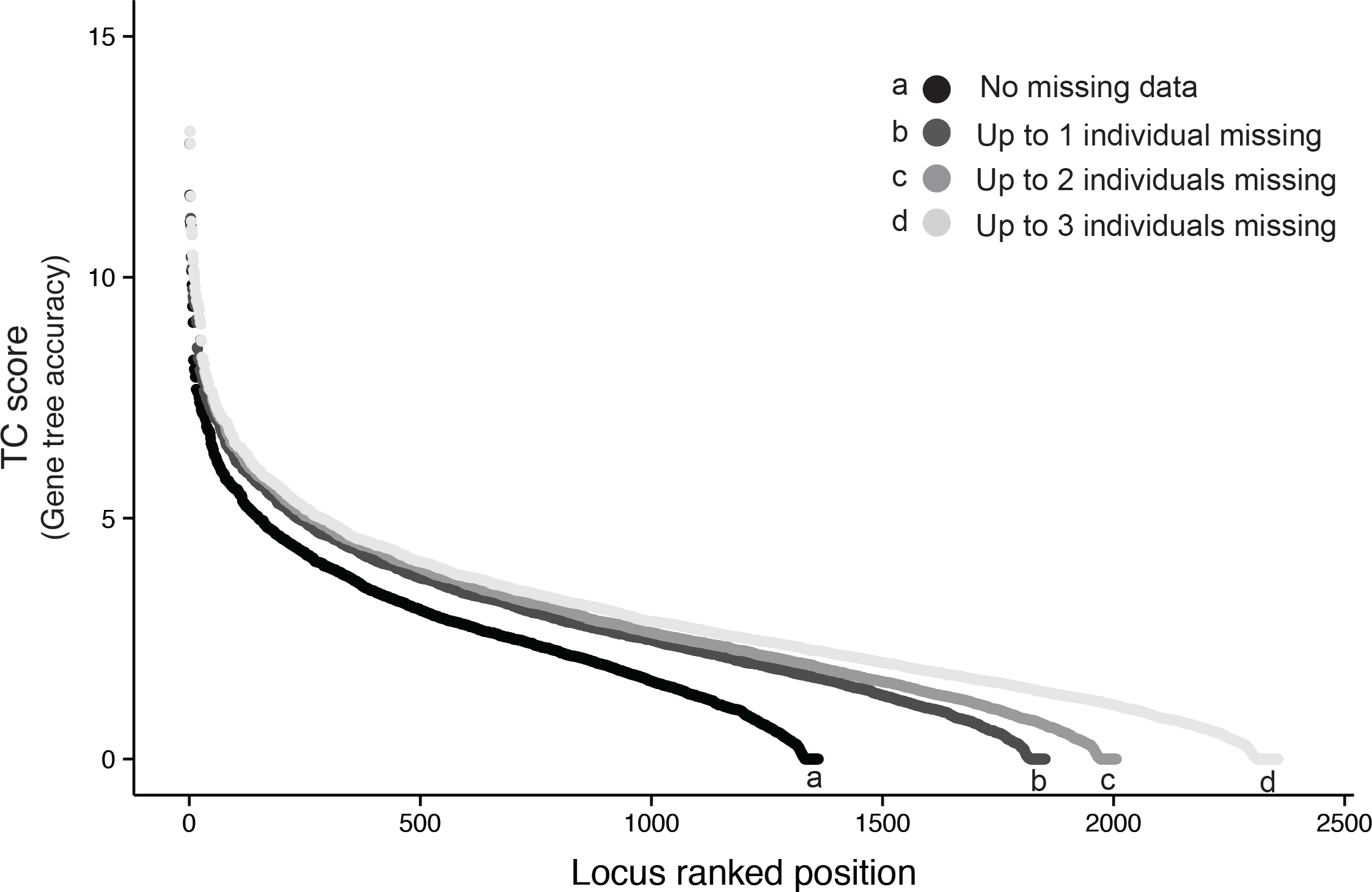
The inferred TC score for each alignment and their ranked position. Different shades (d), correspond to the maximum number of missing species allowed per alignment. By relaxing the criterion for missing species, the total number of recovered loci and the average TC score, gene tree accuracy, for a given number of loci increased.

TC score was positively correlated with locus length (linear model, N = 1361, r^2^ = 0.369, *p* < 0.01; Supplementary Fig. S2), suggesting that longer loci on average result in more accurate gene trees (Liu et al. 2015b). Previous studies have recommended several alternative features that increase the PIC of loci, such as high mutation rate (Lanier et al. 2014; Giarla and Esselstyn 2015), but we expect that locus length is likely the most unbiased surrogate to select for and, as demonstrated here, can significantly increase gene tree accuracy. While aiming for loci with a high mutation rate would induce an increase in the number of parsimony informative characters, it might also bias branch length estimation or result in erroneous inference due to saturation. These results (Supplementary Fig. S2) suggest it might be advantageous to target long loci where possible in sequence-capture studies and we encourage further technical development on this front (e.g. Faircloth et al. 2012; Lemmon et al. 2012). The use of long loci might have some adverse effects if it increases the risk of intra-locus recombination; but simulation experiments suggest that a limited amount of recombination within loci has a relatively modest effect on the inferred species tree (Lanier and Knowles 2012).

### Concatenated and Summary-Coalescent Species Trees

The ML phylogeny (Fig. 3), based on concatenation of the complete dataset (1361 exons, no missing taxa: 522,592 bp), provides a significant gain in resolution of the evolutionary relationships of Australian *Cryptoblepharus* in comparison to the Neighbor-joining (NJ) tree based on allozyme distances (Horner and Adams 2007). As in the allozyme analysis, the Australian representatives of the *Cryptoblepharus* genus are clearly separated in two distinct clades. The few well-supported bipartitions in the allozyme NJ tree are also present in the ML phylogeny, and there are only two nodes in the concatenated exon-capture analysis where the interspecific relationships are poorly resolved (BS < 90). However, the short branch lengths between many nodes in clade B warrant the application of species tree estimation methods that account for deep coalescence.

**Figure 3.**
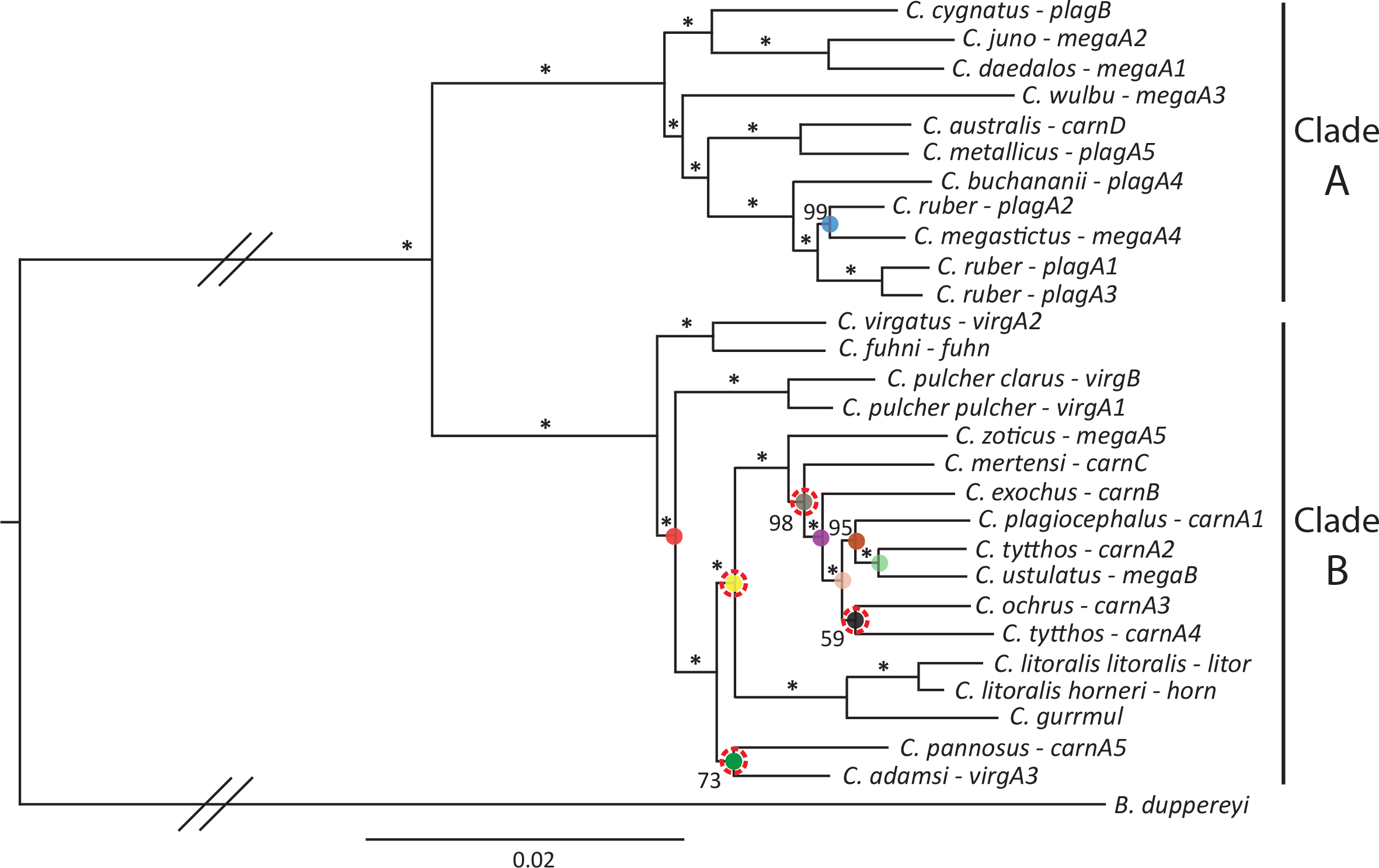
Phylogeny of Australian *Cryptoblepharus* based on a concatenated maximum-likelihood analysis of 1361 exons (complete dataset). Values at internode branches reflect bootstrap support (BS) and an asterisk (*) denotes BS = 100. The nodes annotated with colored orbs, were poorly supported (average BS < 95) using ASTRAL with 80 ranked loci and color scheme used matches across figures (Fig. 4 and 5). Colored nodes were encircled with a dotted red line if the topology differed in the ASTRAL analyses. The current species name and allozyme group (sensu Horner and Adams, 2007) are provided at each tip.

ASTRAL summary-coalescent species trees were inferred based on all loci for both the complete (Supplementary Fig. S3) and incomplete dataset (Fig. 4a). The species tree inferred for each dataset generally agrees with the concatenated ML phylogeny and there are no well-supported differences, except for the placement of *C. pannosus*. In the ASTRAL trees, *C. pannosus* is within the *C. gurrmul* et al. clade rather than a sister to this group, as suggested by the concatenated analysis (Fig. 3). However, the ASTRAL species tree based on the complete dataset is not fully resolved, with multiple unresolved nodes (BS < 95) that were highly supported in the concatenated ML phylogeny. For this species tree, the relationships remain unclear between *C. pulcher pulcher/C. pulcher clarus* and *C. virgatus/C. fuhni, C. zoticus* and *C. mertensi*, and within the clade involving *C. exochus/C. tytthos(carnA4)/C. ochrus/C. plagiocephalus/C. ustulatus/C. tytthos(carnA2)* (Supplementary Fig. S3). In comparison, the ASTRAL species tree based on the incomplete dataset, with 491 additional loci, has better overall support. The polytomy between *C. pulcher pulcher/C. pulcher clarus* and *C. virgatus/C. fuhni* is resolved and is consistent with the concatenated ML tree (Fig. 4a). In addition, the resolution within the clade *C. exochus/C. tytthos(carnA4)/C. ochrus/C. plagiocephalus/C. ustulatus/C. tytthos(CarnA2)* is improved with increased BS for each inferred bipartition (Fig. 4b).

**Figure 4.**
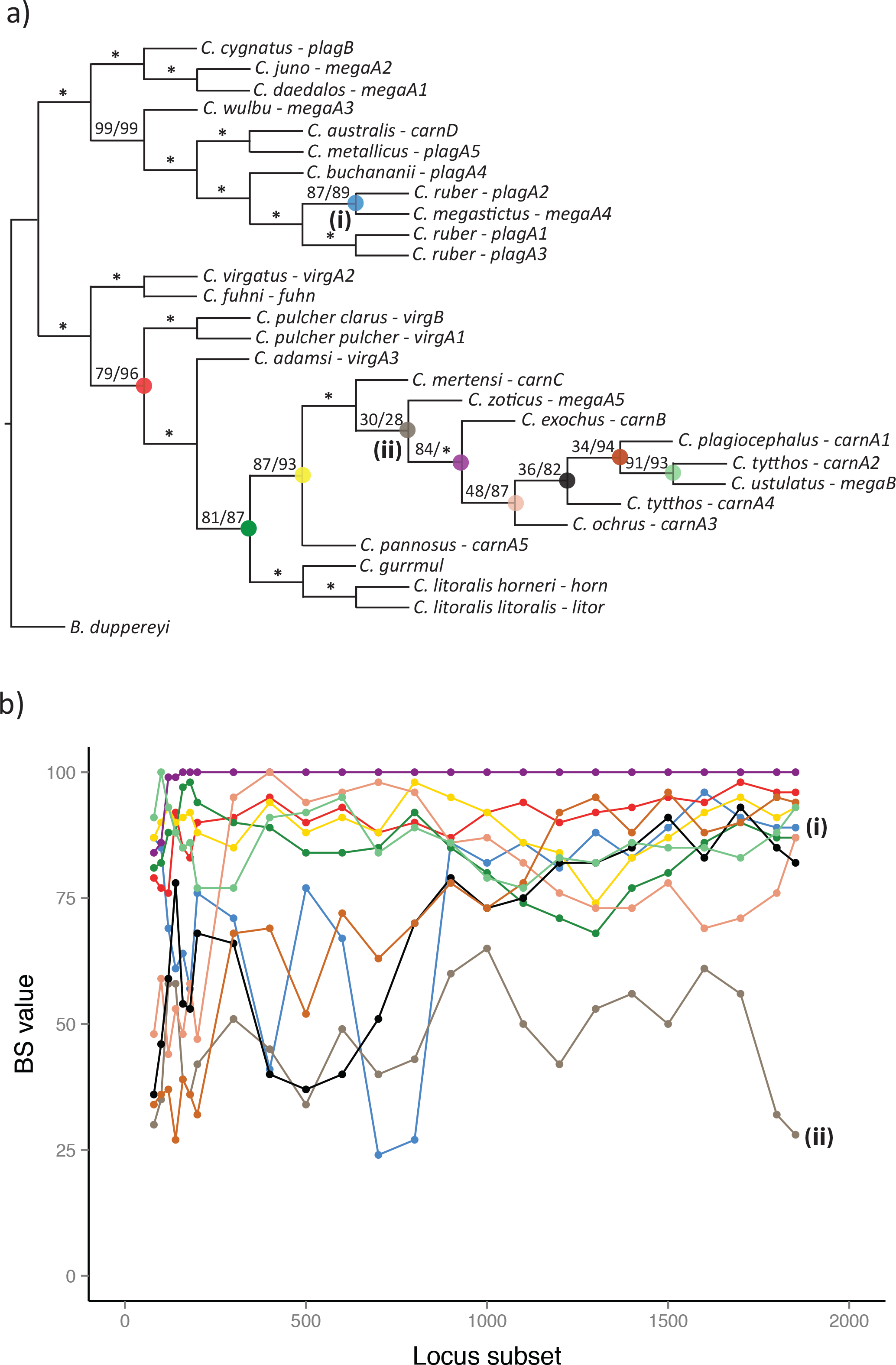
a) Species topology of Australian *Cryptoblepharus*, based on summary-coalescent (ASTRAL) analyses of 80 or 1852 ranked exons (incomplete dataset). The species tree topology changed between initial subsets (20, 40 and 60 loci), but is generally stable from subsets with 80 ranked exons and above. Values at internode branches reflect BS and an asterisk (*) denotes BS = 100. Where two BS values are denoted, the first one is for the species tree based on 80 ranked exons and the second value is for the species tree based on all 1852 exons. The nodes annotated with colored orbs, were poorly supported (average BS < 95) using 80 ranked loci. The current species name and allozyme group (sensu Horner and Adams, 2007) is provided at each tip. b) The change in BS across ranked locus subsets, for each node that was poorly supported (average BS < 95) using 80 ranked loci. The support for two nodes changed erratically, due to potential model violation of the structured coalescent (i.e. introgression) as described in the main text and are labeled (i) and (ii). The color scheme used matches across figures (Fig. 3,4a and 5).

Internode branch lengths in the RAxML phylogeny were positively correlated (N = 6, r^2^ = 0.56, *p* = 0.09, Fig. 5) with average BS values across ASTRAL species trees, suggesting that longer periods between species splits are associated with greater concordance. Each bipartition in the RAxML phylogeny with an internode branch length shorter than 0.0015 (average expected substitutions per site) had an average BS value below 100 in the summary-coalescent analysis. However, the majority of these challenging bipartitions were still strongly supported in the concatenated ML phylogeny (Fig. 3). Even though our ASTRAL species tree based on the incomplete dataset largely agrees with the RAxML phylogeny, caution is warranted regarding the strong support in the ML phylogeny for those bipartitions that are not consistently recovered across species tree inference methods (i.e. placement of *C. pannosus*). Numerous previous studies (e.g. Kubatko and Degnan 2007; Edwards et al. 2007; Knowles 2009; Roch and Steel 2015) have highlighted the potential for inflated BS in phylogenomic size datasets and our results support a similar conclusion. Nonetheless, a concatenated ML phylogeny is computationally cheap in comparison to many other species tree methods and our results demonstrate that, as expected, short branch lengths are indicative for diversification histories where a coalescent-based method is preferred.

**Figure 5.**
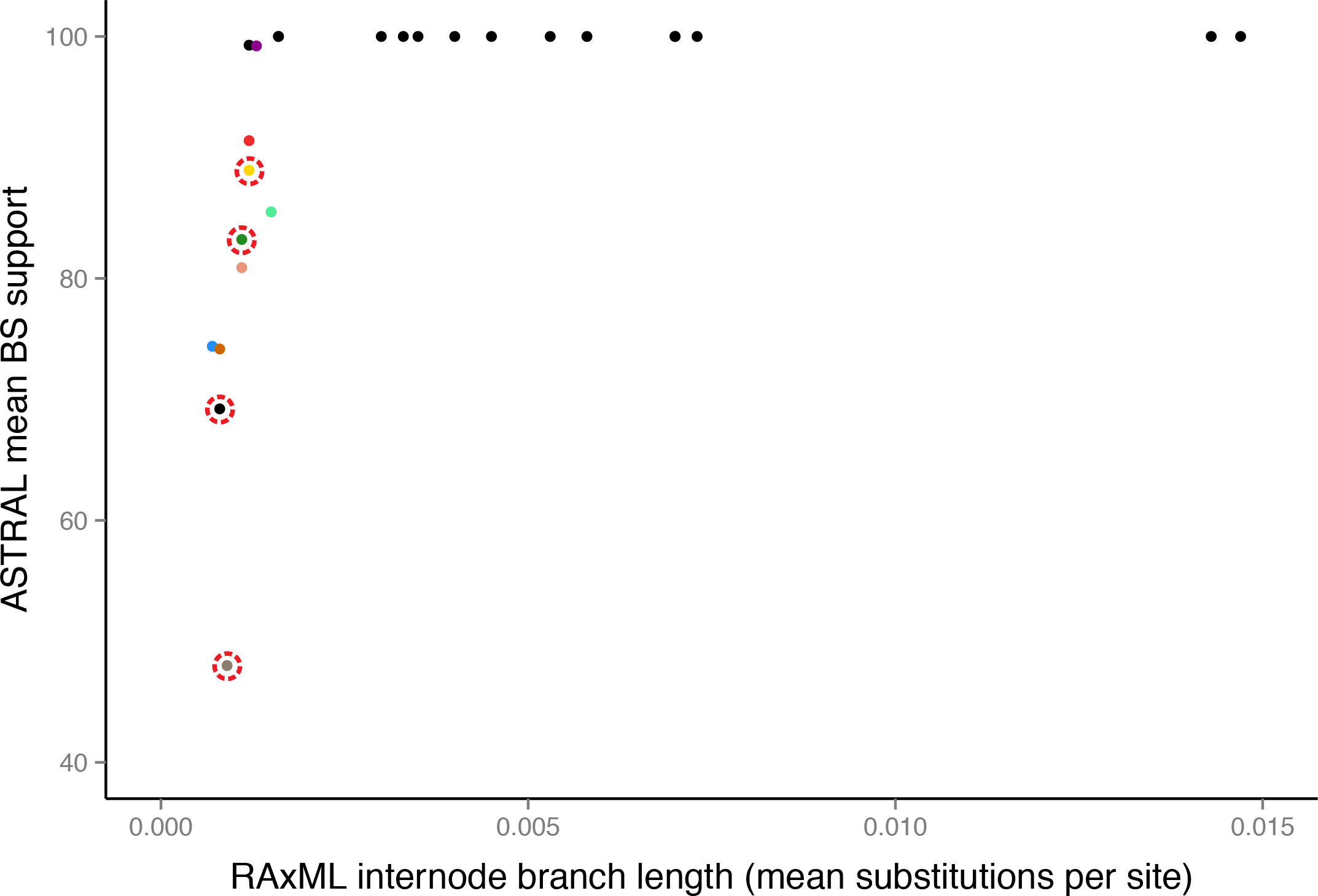
The RAxML internode branch length and mean BS across ranked locus subsets (range 100-1800 loci, incomplete dataset) for each node. Colors match scheme used in other figures (Fig. 3 and 4). Nodes not recovered in the ASTRAL analyses are encircled with a dotted red line.

### Effect of Locus Subsampling by TC score

To evaluate the effect of including gene trees with an increased probability of inaccurate inference on summary-coalescent species tree estimation, we characterized changes in both topology and support. First, we discuss changes in topology with an increase in number of loci while disregarding gene tree resolution. Second, we specifically examine the contrasting patterns of topological change between ranked and random locus subsets of increasing size. Third, in addition to topological variation, we also consider the changes in BS with increasing numbers of ranked loci. Fourth, we discuss whether the addition of low-resolution gene trees introduces phylogenetic noise or, in fact, can increase node support. Finally, we evaluate whether Full-coalescent species tree methods such as *BEAST benefit from the inclusion of loci with high TC scores, and whether joint estimation of gene and species trees lead to improved species tree estimation with similar numbers of loci.

*Topology: number of loci*.— The emergent patterns of topological change with increasing numbers of loci are similar for both the complete (all taxa present; Supplementary Fig. S4) and incomplete dataset (up to one taxa missing; Fig. 6). The RF distance between the species tree for each locus subset and the optimal species tree (i.e. based on all loci) decreases when species trees were inferred based on larger subsets of loci. There was no clear difference if we used the species tree inferred for each previous locus subset, instead of the optimal species tree, for comparison. The decrease in RF distance with additional loci, for both tree comparisons, suggests that topological similarity is not solely induced by an increasing similarity in number of loci between each subset and all loci. If variation in topology was driven by similarity in locus number alone, we would not expect the same decrease in RF distance with additional loci, when comparing the species tree for each subset with the species tree based on the previous locus subset. Even with the observed decrease in RF distance with a larger number of loci, some polytomies remain unresolved. The RF distance between trees therefore does not reduce to zero; even when large numbers of loci are included (Fig. 6). This remaining variation in RF distance depends on how polytomies (i.e. poor support) are ‘arbitrarily’ resolved in each tree, to represent a bifurcating tree. The fluctuation in RF scores is larger for the complete dataset (Supplementary Fig. S4), than for the incomplete dataset (Fig. 6), since the species tree based on the incomplete dataset is more confidently resolved (Fig. 4).

**Figure 6.**
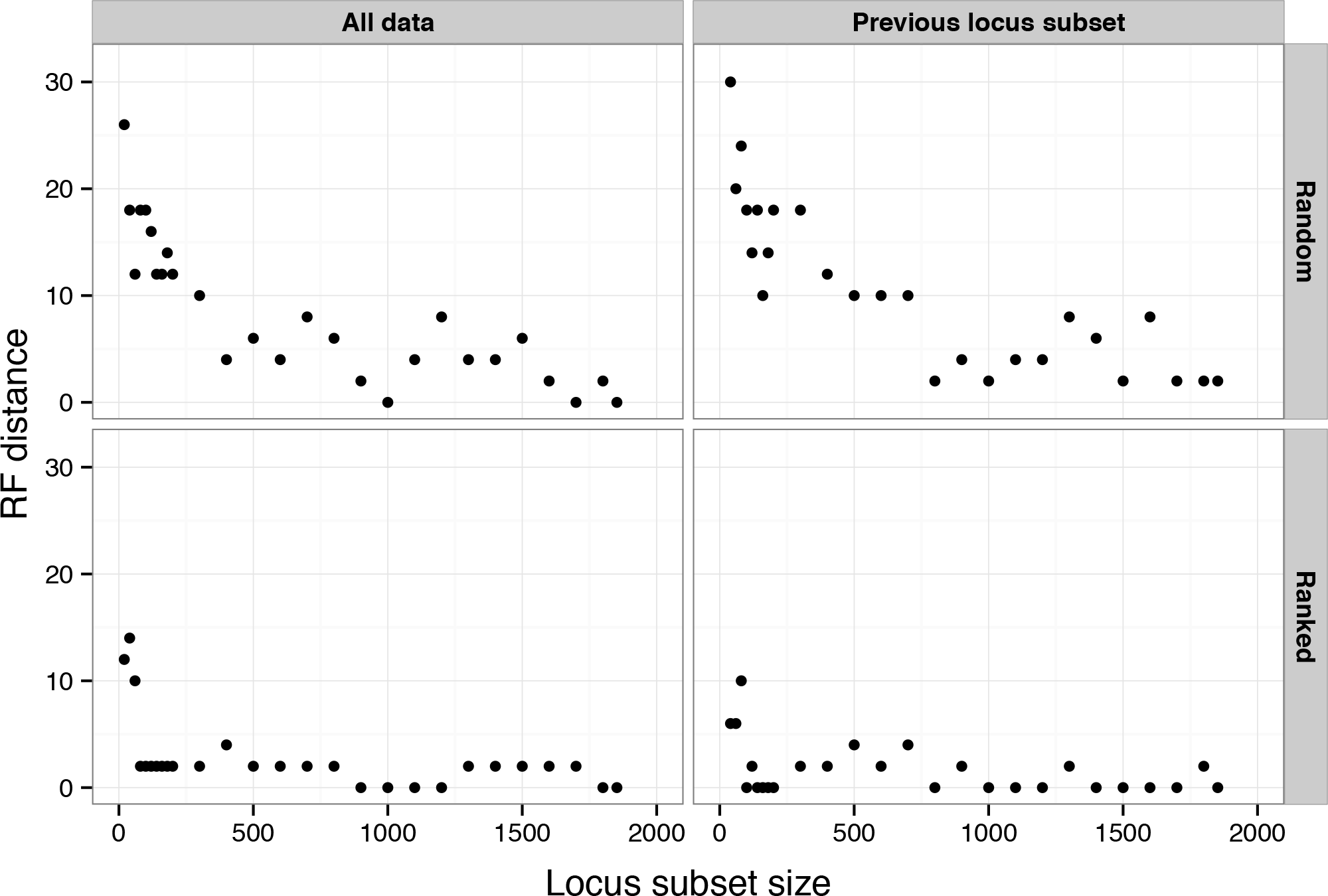
The Robinson-Foulds distance between the inferred ASTRAL species tree for each locus subset (incomplete dataset) and the ASTRAL species trees based on 1852 exons (left) or the previous locus subset (right). Subsets of loci were either chosen randomly (top) or ranked (bottom) by Tree Certainty score. The addition of loci with poor gene tree resolution does not increase topological discordance and limited numbers of ranked loci already converge on the most optimal topology given the dataset.

*Topology: ranked vs. random gene trees*.— When inferring species trees based on ranked subsets of loci, the most optimal topology given the dataset is inferred with fewer loci than if equal sized subsets were picked randomly (Supplementary Fig. S4, Fig. 6). For the incomplete dataset, the inferred species tree based on the 80 loci with the highest TC scores is almost identical in topology to the species tree based on all 1852 loci (Fig. 4a). These trees only differ in how the polytomy involving *C. zoticus* and *C. mertensi* is resolved. The difference between ranked and randomly selected loci dissipates when the size of the locus subset increases, potentially due to the increased likelihood that the randomly selected subsets also contain high-resolution gene trees. In line with previous simulation studies that have evaluated summary-coalescent methods (Lanier et al. 2014; Liu et al. 2015b), high-resolution gene trees seem to have a stronger effect on the inferred topology than low-resolution gene trees. This is further illustrated when comparing the species trees based on small subsets of ranked loci for the incomplete dataset, with the species trees inferred for the complete dataset. The RF distances between trees in the complete dataset are higher, even for reasonably large subsets of loci, than when comparing between trees based on small subsets of ranked loci for the incomplete dataset (Fig.6, Supplementary Fig. S4). This difference is likely due to the larger number of high-resolution gene trees in the incomplete dataset (Fig. 2).

*Bootstrap support*.— Due to the minimal differences in topology between subsets of ranked loci (incomplete dataset), we evaluated the change in BS values for each bipartition (Fig. 4b) across species trees based on subsets of increasing size. Whereas topological changes are limited, the BS values for some nodes change substantially with increasing numbers of loci. Nodes that are well supported (BS = 100) remain high, but different patterns emerge for the more challenging nodes. For the nodes that had a BS value < 90 in the species tree based on 80 ranked loci (Fig. 4a), the accuracy increases on average with additional loci (Fig. 4b). But whereas BS increases coherently for some difficult nodes (e.g. the placement of the *C. pulcher* clade), BS estimates for other nodes fluctuate considerably with increasing size of locus subsets. Most notably, the BS for the polytomy involving *C. zoticus* and *C. mertensi* on average fluctuates from 40 a 60 and ultimately, remains unresolved. Secondly, the BS values for the node involving *C. ruber(plagA2)* and *C. megastictus* change markedly between subsets, up to 1000 ranked loci, and then steadily increases for larger consecutive subsets. Interestingly the taxa for which BS values vary erratically with increasing subset size also match observations of mitochondrial introgression between distinct ecomorphs; specifically in these two species pairs (*C. ruber/C. megastictus, C. zoticus/C. mertensi;* Blom et al., *in prep.*). These results suggest possible nuclear introgression and violation of the structured coalescent model. Most coalescent-based species tree methods assume that incongruence is solely due to incomplete lineage sorting, but introgression could result in similar patterns of topological discordance (Kutschera et al. 2014; Nater et al. 2015). By quantifying patterns of change in topology or support across locus subsets, such underlying signals can be distilled, inspected for biological relevance and potentially studied in a coalescent framework that also aims to model alternative sources of gene tree - species tree incongruence (i.e. a phylogenetic network approach (Yu et al. 2014).

*Phylogenetic noise*.— The addition of gene trees with low TC scores increases the probability of including gene trees with stochastic estimation error. However, this did not result in any well-supported topological changes (Fig. 4a). For both the complete and incomplete dataset, a number of nodes remain unresolved and some topological variance between trees was generated depending on how such uncertain relationships were (arbitrarily) resolved. The remaining variation in RF distances between trees does not imply that the addition of low-resolution loci reduces the accuracy of the species tree estimate since we would expect that this would have a similar effect on well-supported nodes. The topological uncertainty driving the persistent variability in RF distances between trees based on large locus subsets only occurs at nodes that were poorly supported throughout the analyses.

Rather than introducing phylogenetic noise (Liu et al. 2015b), our results suggest that the inclusion of low-resolution gene trees increases the consistency of phylogenetic inference with average BS values increasing across several of the most challenging bipartitions (Fig. 4b). Previous simulation studies (Leache and Rannala 2011; Mirarab et al. 2014a) have highlighted that for most relationships, barring the ‘anomaly zone’ (Liu and Edwards 2009; Huang and Knowles 2009), coalescent-based approaches will infer the correct phylogenetic history with sufficient numbers of independent loci. Even though most of these analyses assume a homogeneous distribution of PIC across genes, our results suggest that even relatively uninformative loci still improved BS across some of the most challenging bipartitions (Fig. 4b). This was surprising to us and we encourage future simulation studies to further examine this empirical observation. We do not expect that this is an artifact of the method used since other nodes, such as the placement of *C. zoticus/C. mertensi*, remain unresolved; regardless of the number of gene trees included.

*Full-coalescent analyses with ranked loci*.— Lastly, we evaluated whether a small subset of the most informative loci could potentially be sufficient to infer the optimal species tree using a Full-coalescent species tree analysis. Although the joint estimation of gene and species tree is generally preferred over summary-coalescent methods (Leache and Rannala 2011; Ogilvie et al. 2016), we did not infer a more accurate species tree using *BEAST (Supplementary Fig. S5). The *BEAST species trees differed from the optimal species tree to a similar extent as ASTRAL trees based on ranked loci subsets of similar size (Supplementary Fig. S5). Furthermore, *BEAST trees based on similar sized (20 or 30 loci) random subsets of loci with the highest TC scores vary substantially (Supplementary Fig. S5). Currently, we cannot determine with certainty whether the differences between the *BEAST trees and the ASTRAL tree based on all loci, occur due to methodological differences between full and summary-coalescent analyses, or if it is because *BEAST can only be run with a much smaller number of loci. Though the discordance among the *BEAST trees suggests the latter. However we expect this to be a fruitful area of research in the future, with improvements in the implementation of Full-coalescent species tree analyses.

## Conclusion

Species trees are commonly estimated based on gene trees. It is well known that those gene trees can be different from the species tree due to a variety of biological processes. For example, incomplete lineage sorting can be explicitly incorporated into species tree analyses, using coalescent-based methods. However, gene trees can also differ from the underlying species tree due to estimation error and unlike most simulation analyses, empirical phylogenomic studies often generate gene trees that widely vary in resolution. Here we quantified the impact of stochastic gene tree estimation error on summary-coalescent species tree inference by screening loci based on gene tree resolution prior to species tree estimation. Our results indicate that longer loci yield higher resolution gene trees, that a relatively small number (~80, incomplete dataset) of high resolution gene trees can already converge on the optimal topology given the dataset and that convergence on this topology occurs with fewer loci if gene trees are first ordered by resolution. Moreover, the addition of low-resolution gene trees did not introduce phylogenetic noise (Liu et al. 2015b) and in fact increased support for several challenging nodes. These empirical findings highlight the importance of gene tree resolution for species tree inference and are in line with previous simulation studies (Lanier et al. 2014; Mirarab et al. 2014a; Liu et al. 2015b).

Unlike most simulations, empirical phylogenomic datasets are highly heterogeneous in terms of phylogenetic information content. Rather than treating each sequenced locus as equal, our study suggests that by characterizing the heterogeneity in gene tree resolution and ranking them accordingly, we can account for stochastic gene tree estimation error when using summary-coalescent methods. Although it remains unclear to what extent our empirical findings can be generalized across taxa with different diversification histories, we have presented a conceptual approach that can be applied without difficulty in any other phylogenomic study where sequenced loci vary in PIC. By using the TC score of gene trees as a proxy for gene tree resolution, we have demonstrated how the ranking of loci by gene tree resolution prior to summary-coalescent species tree estimation can yield valuable insights. In addition to providing the means to evaluate the effect of including gene trees with an increased probability of inaccurate inference, it also enabled us to assess the importance of gene tree resolution in general and to identify characteristics of loci that can predict the accuracy of gene tree estimates (i.e. length). Thus our results strongly suggest that the evaluation of gene tree resolution is an informative and relevant practice that can greatly benefit empiricists.

Whereas summary-coalescent methods might prove effective in estimating a tree topology, Full-coalescent sequence based methods can simultaneously infer a topology and branch lengths, and are therefore ultimately more appropriate for many purposes. A question that remains unexplored is to what extent a hybrid approach that incorporates both summary- and full-coalescent analyses can improve the computational efficiency and accuracy of species tree inference? Our results show that the majority of the *Cryptoblepharus* topology was unambiguously supported across most subsets of loci and that a small subset of informative loci already inferred the most optimal topology. If computationally efficient approaches such as summary-coalescent methods can resolve ‘easy’ nodes confidently, the search space of possible trees could be reduced significantly for a Full-coalescent method. It remains unknown whether these implementations would reduce the computational challenge of full-coalescent species tree estimation and allow for the inclusion of more loci, but it merits further investigation. In conclusion, our findings suggest there is much scope for future theoretical and empirical research into the best approaches for estimating species trees with large phylogenetic datasets. Such studies should explicitly consider the heterogeneity in phylogenetic signal among loci and how this translates into an accurate inference of the underlying species tree.

## Acknowledgements

We thank Paul Horner and Mark Adams who generously shared the morphological and allozyme data, which were beneficial to target appropriate representatives of each species. Ke Bi and Sonal Singhal provided guidance with both laboratory and bioinformatic procedures. M.P.K.B greatly benefited from the Computational Molecular Evolution EMBO workshop. We thank Huw Ogilvie and Robert Lanfear for insightful discussions regarding species tree inference analysis approaches. We thank the Australian Biological Tissue Collection at the South Australian Museum (Stephen Donnellan), the Western Australian Museum (Paul Doughty), the Queensland Museum (Jessica Worthington Wilmer) and the Museum and Art Gallery of the Northern Territory (Stephen Richards) for access to tissues and specimens.

